## Supplement 1 for "Failure of the brain glucagon-like peptide-1-mediated control of intestinal redox homeostasis in a rat model of sporadic Alzheimer’s disease"

Table 1. Plasma SOD – main effects model

| contrast | estimate | SE | df | lower.CL | upper.CL | t.ratio | p.value |
|---|---|---|---|---|---|---|---|
| CTR - STZ | 2.5185 | 0.5706 | 36 | 1.3613 | 3.6757 | 4.4140 | 0.0001 |
| (Ex-9) - SAL | -0.6483 | 0.5706 | 36 | -1.8054 | 0.5089 | -1.1362 | 0.2634 |

Table 2. Plasma SOD – treatment interaction model

| contrast | estimate | SE | df | lower.CL | upper.CL | t.ratio | p.value |
|---|---|---|---|---|---|---|---|
| (CTR Ex-9) - (STZ Ex-9) | 1.3197 | 0.7800 | 35 | -0.7838 | 3.4232 | 1.6920 | 0.3430 |
| (CTR Ex-9) - CTR SAL | -1.7840 | 0.7592 | 35 | -3.8314 | 0.2634 | -2.3500 | 0.1061 |
| (CTR Ex-9) - STZ SAL | 1.8703 | 0.7592 | 35 | -0.1771 | 3.9176 | 2.4636 | 0.0837 |
| (STZ Ex-9) - CTR SAL | -3.1037 | 0.7800 | 35 | -5.2071 | -1.0002 | -3.9793 | 0.0018 |
| (STZ Ex-9) - STZ SAL | 0.5506 | 0.7800 | 35 | -1.5529 | 2.6540 | 0.7059 | 0.8941 |
| CTR SAL - STZ SAL | 3.6542 | 0.7592 | 35 | 1.6069 | 5.7016 | 4.8136 | 0.0002 |

Table 3. plasma TBARS – main effects model

| contrast | estimate | SE | df | lower.CL | upper.CL | t.ratio | p.value |
|---|---|---|---|---|---|---|---|

| | | | | | | | |
|---|---|---|---|---|---|---|---|
| CTR - STZ | -0.0082 | 0.0051 | 36 | -0.0186 | 0.0022 | -1.6004 | 0.1183 |
| (Ex-9) - SAL | -0.0140 | 0.0051 | 36 | -0.0244 | -0.0035 | -2.7201 | 0.0100 |

Table 4. Plasma TBARS – interaction model

| contrast | estimate | SE | df | lower.CL | upper.CL | t.ratio | p.value |
|---|---|---|---|---|---|---|---|
| (CTR Ex-9) - (STZ Ex-9) | -0.0083 | 0.0075 | 35 | -0.0284 | 0.0118 | -1.1143 | 0.6833 |
| (CTR Ex-9) - CTR SAL | -0.0140 | 0.0073 | 35 | -0.0336 | 0.0055 | -1.9353 | 0.2322 |
| (CTR Ex-9) - STZ SAL | -0.0222 | 0.0073 | 35 | -0.0417 | -0.0026 | -3.0528 | 0.0214 |
| (STZ Ex-9) - CTR SAL | -0.0057 | 0.0075 | 35 | -0.0258 | 0.0144 | -0.7694 | 0.8677 |
| (STZ Ex-9) - STZ SAL | -0.0138 | 0.0075 | 35 | -0.0340 | 0.0063 | -1.8571 | 0.2648 |
| CTR SAL - STZ SAL | -0.0081 | 0.0073 | 35 | -0.0277 | 0.0115 | -1.1175 | 0.6813 |

Table 5. Plasma NRP – main effects model

| contrast | estimate | SE | df | lower.CL | upper.CL | t.ratio | p.value |
|---|---|---|---|---|---|---|---|
| CTR - STZ | 2906783 | 877107.3 | 36 | 1127926 | 4685639 | 3.3141 | 0.0021 |
| (Ex-9) - SAL | -3295336 | 877107.3 | 36 | -5074192 | -1516480 | -3.7571 | 0.0006 |

Table 6. Plasma NRP – interaction model

| contrast | estimate | SE | df | lower.CL | upper.CL | t.ratio | p.value |
|---|---|---|---|---|---|---|---|
| (CTR Ex-9) - (STZ Ex-9) | 2296262.9 | 1267209 | 35 | -1121275.9 | 5713801.6 | 1.8121 | 0.2849 |
| (CTR Ex-9) - CTR SAL | -3873723.3 | 1233410 | 35 | -7200111.2 | -547335.4 | -3.1407 | 0.0172 |
| (CTR Ex-9) - STZ SAL | -388553.6 | 1233410 | 35 | -3714941.5 | 2937834.3 | -0.3150 | 0.9890 |
| (STZ Ex-9) - CTR SAL | -6169986.2 | 1267209 | 35 | -9587524.9 | -2752447.4 | -4.8690 | 0.0001 |
| (STZ Ex-9) - STZ SAL | -2684816.5 | 1267209 | 35 | -6102355.2 | 732722.3 | -2.1187 | 0.1671 |
| CTR SAL - STZ SAL | 3485169.7 | 1233410 | 35 | 158781.8 | 6811557.6 | 2.8256 | 0.0371 |

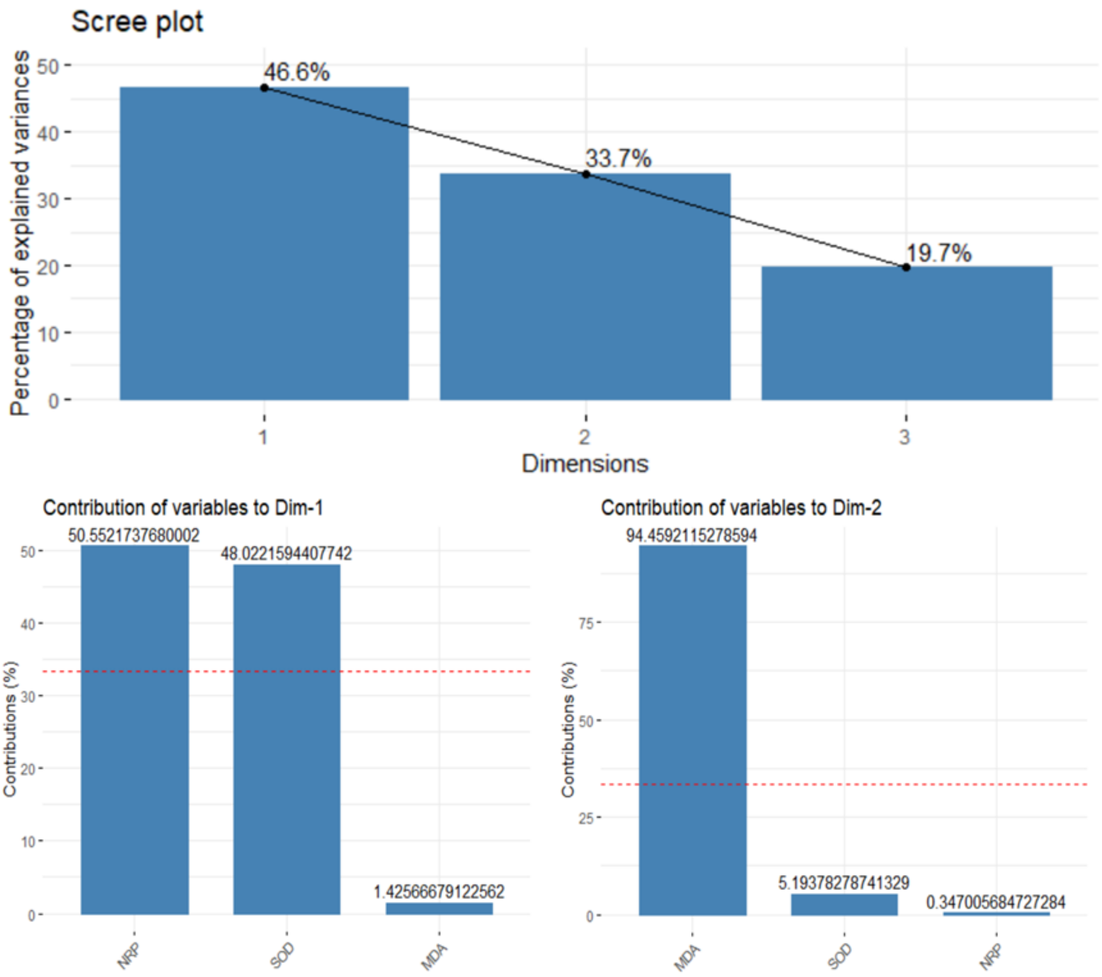


Fig 1. Scree plot and contribution of individual variables to dimensions for plasma oxidative stress PCA.

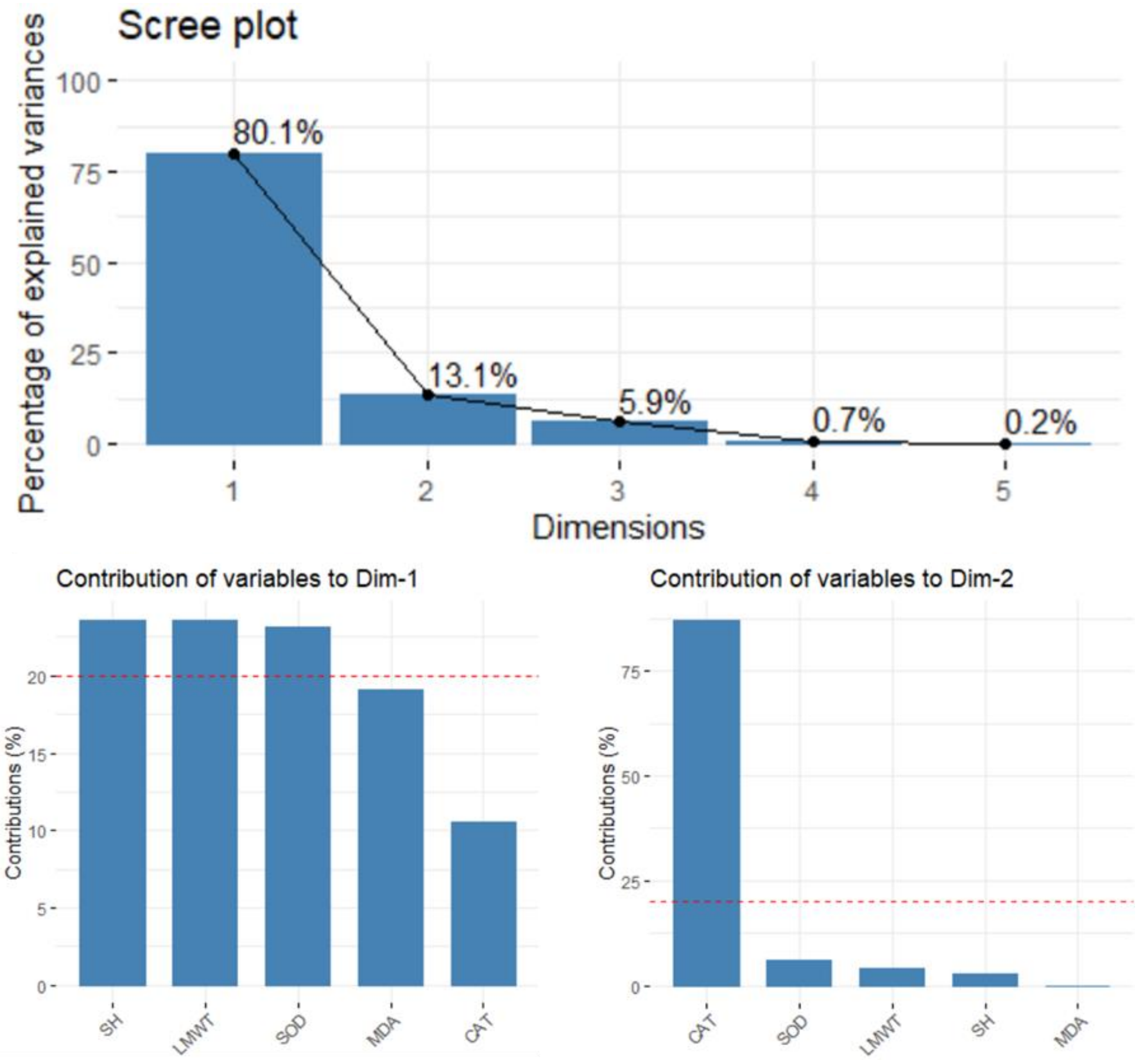


Fig 2. Scree plot and contribution of individual variables to dimensions for duodenum oxidative stress PCA.

Table 7. Duodenum oxidative stress PCA position relative to Dim 1 – main effects model

| contrast | estimate | SE | df | lower.CL | upper.CL | t.ratio | p.value |
|---|---|---|---|---|---|---|---|
| CTR - STZ | -0.7609 | 0.931 | 16 | -2.7345 | 1.2128 | -0.8172 | 0.4258 |
| (Ex-9) - SAL | 0.9501 | 0.931 | 16 | -1.0236 | 2.9238 | 1.0205 | 0.3227 |

Table 8. Duodenum oxidative stress PCA position relative to Dim 1 – interaction model

| contrast | estimate | SE | df | lower.CL | upper.CL | t.ratio | p.value |
|---|---|---|---|---|---|---|---|
| (CTR Ex-9) - (STZ Ex-9) | 1.1843 | 1.2199 | 15 | -1.4159 | 3.7846 | 0.9708 | 0.3470 |
| (CTR Ex-9) - CTR SAL | 2.6792 | 1.1502 | 15 | 0.2276 | 5.1307 | 2.3293 | 0.0342 |
| (CTR Ex-9) - STZ SAL | 0.1892 | 1.1502 | 15 | -2.2623 | 2.6408 | 0.1645 | 0.8715 |
| (STZ Ex-9) - CTR SAL | 1.4948 | 1.2199 | 15 | -1.1054 | 4.0951 | 1.2253 | 0.2394 |
| (STZ Ex-9) - STZ SAL | -0.9951 | 1.2199 | 15 | -3.5954 | 1.6051 | -0.8157 | 0.4274 |
| CTR SAL - STZ SAL | -2.4899 | 1.1502 | 15 | -4.9415 | -0.0384 | -2.1648 | 0.0469 |

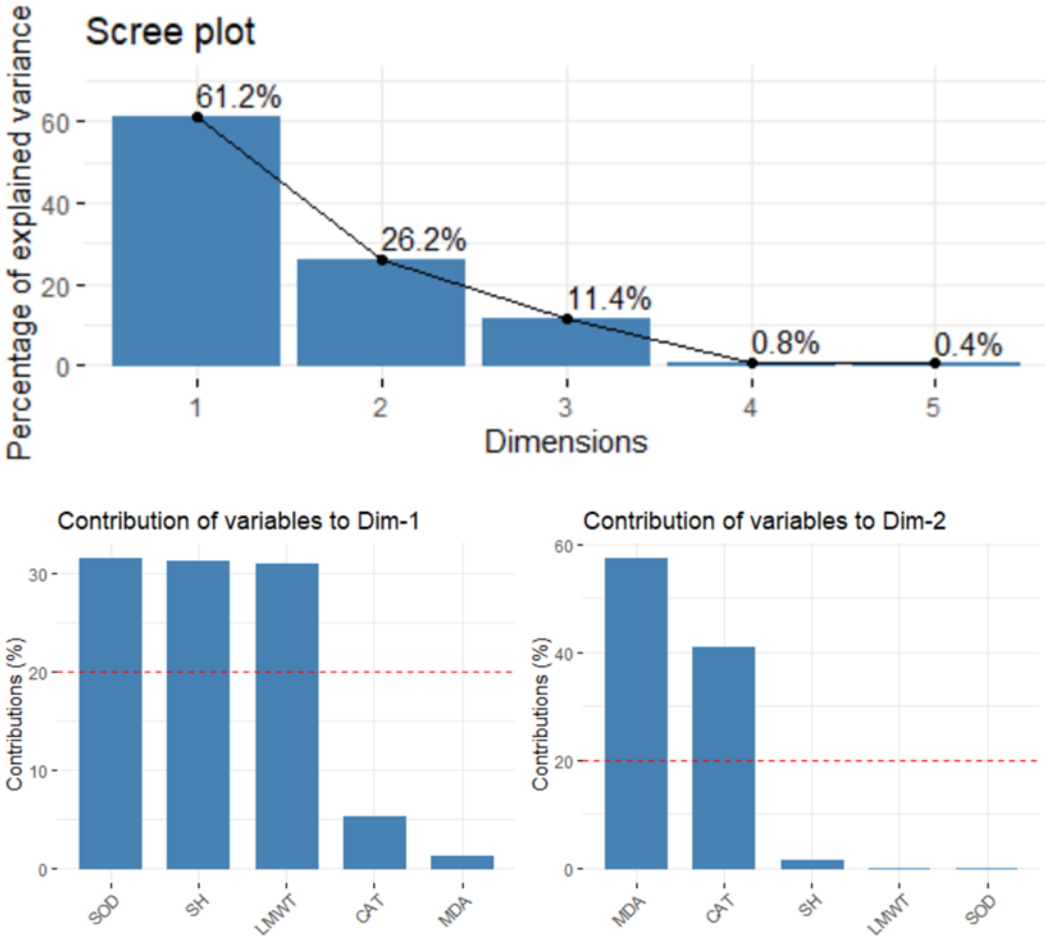


Fig 2. Scree plot and contribution of individual variables to dimensions for ileum oxidative stress PCA.

Table 9. Ileum oxidative stress PCA position relative to Dim 1 – main effects model

| contrast | estimate | SE | df | lower.CL | upper.CL | t.ratio | p.value |
|---|---|---|---|---|---|---|---|
| CTR - STZ | 0.6091 | 0.7543 | 20 | -0.9643 | 2.1825 | 0.8075 | 0.4289 |
| (Ex-9) - SAL | -0.0920 | 0.7543 | 20 | -1.6654 | 1.4814 | -0.1220 | 0.9041 |

Table 10. Ileum oxidative stress PCA position relative to Dim 1 – interaction model

| contrast | estimate | SE | df | lower.CL | upper.CL | t.ratio | p.value |
|---|---|---|---|---|---|---|---|

| | | | | | | | |
|---|---|---|---|---|---|---|---|
| (CTR Ex-9) - (STZ Ex-9) | 0.8147 | 1.0671 | 19 | -1.4187 | 3.0482 | 0.7635 | 0.4545 |
| (CTR Ex-9) - CTR SAL | 0.1342 | 1.1192 | 19 | -2.2082 | 2.4766 | 0.1199 | 0.9058 |
| (CTR Ex-9) - STZ SAL | 0.5171 | 1.0671 | 19 | -1.7163 | 2.7505 | 0.4846 | 0.6335 |
| (STZ Ex-9) - CTR SAL | -0.6805 | 1.1192 | 19 | -3.0230 | 1.6619 | -0.6081 | 0.5503 |
| (STZ Ex-9) - STZ SAL | -0.2976 | 1.0671 | 19 | -2.5311 | 1.9358 | -0.2789 | 0.7833 |
| CTR SAL - STZ SAL | 0.3829 | 1.1192 | 19 | -1.9596 | 2.7253 | 0.3421 | 0.7360 |

Table 11. Duodenum TBARS – main effects model

| contrast | estimate | SE | df | lower.CL | upper.CL | t.ratio | p.value |
|---|---|---|---|---|---|---|---|
| CTR - STZ | 2.1665 | 16.8437 | 16 | -33.5407 | 37.8736 | 0.1286 | 0.8993 |
| (Ex-9) - SAL | 5.9182 | 16.8437 | 16 | -29.7889 | 41.6254 | 0.3514 | 0.7299 |

Table 12. Duodenum TBARS – interaction model

| contrast | estimate | SE | df | lower.CL | upper.CL | t.ratio | p.value |
|---|---|---|---|---|---|---|---|
| (CTR Ex-9) - (STZ Ex-9) | 50.8383 | 18.5679 | 15 | 11.2617 | 90.4149 | 2.7380 | 0.0153 |
| (CTR Ex-9) - CTR SAL | 49.1821 | 17.5060 | 15 | 11.8689 | 86.4953 | 2.8094 | 0.0132 |
| (CTR Ex-9) - STZ SAL | 8.0847 | 17.5060 | 15 | -29.2285 | 45.3979 | 0.4618 | 0.6508 |
| (STZ Ex-9) - CTR SAL | -1.6562 | 18.5679 | 15 | -41.2328 | 37.9204 | -0.0892 | 0.9301 |
| (STZ Ex-9) - STZ SAL | -42.7536 | 18.5679 | 15 | -82.3302 | -3.1770 | -2.3026 | 0.0360 |
| CTR SAL - STZ SAL | -41.0974 | 17.5060 | 15 | -78.4106 | -3.7842 | -2.3476 | 0.0330 |

Table 13. Duodenum LMWT – main effects model

| contrast | estimate | SE | df | lower.CL | upper.CL | t.ratio | p.value |
|---|---|---|---|---|---|---|---|
| CTR - STZ | 3.5732 | 2.8087 | 16 | -2.3810 | 9.5274 | 1.2722 | 0.2215 |
| (Ex-9) - SAL | -4.0849 | 2.8087 | 16 | -10.0391 | 1.8693 | -1.4544 | 0.1652 |

Table 14. Duodenum LMWT – interaction model

| contrast | estimate | SE | df | lower.CL | upper.CL | t.ratio | p.value |
|---|---|---|---|---|---|---|---|
| (CTR Ex-9) - (STZ Ex-9) | -1.6827 | 3.7951 | 15 | -9.7717 | 6.4064 | -0.4434 | 0.6638 |

| | | | | | | | |
|---|---|---|---|---|---|---|---|
| (CTR Ex-9) - CTR SAL | -8.7568 | 3.5780 | 15 | -16.3832 | -1.1304 | -2.4474 | 0.0272 |
| (CTR Ex-9) - STZ SAL | -0.5117 | 3.5780 | 15 | -8.1381 | 7.1148 | -0.1430 | 0.8882 |
| (STZ Ex-9) - CTR SAL | -7.0741 | 3.7951 | 15 | -15.1632 | 1.0149 | -1.8640 | 0.0820 |
| (STZ Ex-9) - STZ SAL | 1.1710 | 3.7951 | 15 | -6.9180 | 9.2601 | 0.3086 | 0.7619 |
| CTR SAL - STZ SAL | 8.2451 | 3.5780 | 15 | 0.6187 | 15.8715 | 2.3044 | 0.0359 |

Table 15. Duodenum SH – main effects model

| contrast | estimate | SE | df | lower.CL | upper.CL | t.ratio | p.value |
|---|---|---|---|---|---|---|---|
| CTR - STZ | 2.3588 | 2.1714 | 16 | -2.2445 | 6.9620 | 1.0863 | 0.2935 |
| (Ex-9) - SAL | -2.9370 | 2.1714 | 16 | -7.5402 | 1.6663 | -1.3526 | 0.1950 |

Table 16. Duodenum SH – interaction model

| contrast | estimate | SE | df | lower.CL | upper.CL | t.ratio | p.value |
|---|---|---|---|---|---|---|---|
| (CTR Ex-9) - (STZ Ex-9) | -1.2986 | 3.0005 | 15 | -7.6940 | 5.0969 | -0.4328 | 0.6713 |
| (CTR Ex-9) - CTR SAL | -6.1879 | 2.8289 | 15 | -12.2176 | -0.1582 | -2.1874 | 0.0450 |

| | | | | | | | |
|---|---|---|---|---|---|---|---|
| (CTR Ex-9) - STZ SAL | -0.5782 | 2.8289 | 15 | -6.6079 | 5.4515 | -0.2044 | 0.8408 |
| (STZ Ex-9) - CTR SAL | -4.8893 | 3.0005 | 15 | -11.2848 | 1.5061 | -1.6295 | 0.1240 |
| (STZ Ex-9) - STZ SAL | 0.7204 | 3.0005 | 15 | -5.6751 | 7.1158 | 0.2401 | 0.8135 |
| CTR SAL - STZ SAL | 5.6097 | 2.8289 | 15 | -0.4200 | 11.6394 | 1.9830 | 0.0660 |

Table 17. Duodenum CAT – main effects model

| contrast | ratio | SE | df | lower.CL | upper.CL | t.ratio | p.value |
|---|---|---|---|---|---|---|---|
| CTR / STZ | 0.9714 | 0.5584 | 16 | 0.2872 | 3.2856 | -0.0504 | 0.9604 |
| (Ex-9) / SAL | 0.7011 | 0.4030 | 16 | 0.2073 | 2.3713 | -0.6178 | 0.5454 |

Table 18. Duodenum CAT – interaction model

| contrast | ratio | SE | df | lower.CL | upper.CL | t.ratio | p.value |
|---|---|---|---|---|---|---|---|
| (CTR Ex-9) / (STZ Ex-9) | 3.8914 | 2.7691 | 15 | 0.8538 | 17.7346 | 1.9094 | 0.0755 |
| (CTR Ex-9) / CTR SAL | 2.4072 | 1.6150 | 15 | 0.5760 | 10.0590 | 1.3093 | 0.2101 |
| (CTR Ex-9) / STZ SAL | 0.6811 | 0.4570 | 15 | 0.1630 | 2.8461 | -0.5724 | 0.5755 |

| | | | | | | | |
|---|---|---|---|---|---|---|---|
| (STZ Ex-9) / CTR SAL | 0.6186 | 0.4402 | 15 | 0.1357 | 2.8192 | -0.6750 | 0.5100 |
| (STZ Ex-9) / STZ SAL | 0.1750 | 0.1246 | 15 | 0.0384 | 0.7977 | -2.4491 | 0.0271 |
| CTR SAL / STZ SAL | 0.2829 | 0.1898 | 15 | 0.0677 | 1.1824 | -1.8818 | 0.0794 |

Table 19. Duodenum SOD – main effects model

| contrast | estimate | SE | df | lower.CL | upper.CL | t.ratio | p.value |
|---|---|---|---|---|---|---|---|
| CTR - STZ | 0.0207 | 0.0236 | 16 | -0.0294 | 0.0708 | 0.8745 | 0.3948 |
| (Ex-9) - SAL | -0.0379 | 0.0236 | 16 | -0.0880 | 0.0122 | -1.6018 | 0.1288 |

Table 19. Duodenum SOD – interaction model

| contrast | estimate | SE | df | lower.CL | upper.CL | t.ratio | p.value |
|---|---|---|---|---|---|---|---|
| (CTR Ex-9) - (STZ Ex-9) | -0.0158 | 0.0331 | 15 | -0.0864 | 0.0549 | -0.4760 | 0.6409 |
| (CTR Ex-9) - CTR SAL | -0.0703 | 0.0313 | 15 | -0.1369 | -0.0036 | -2.2480 | 0.0400 |
| (CTR Ex-9) - STZ SAL | -0.0172 | 0.0313 | 15 | -0.0838 | 0.0494 | -0.5500 | 0.5904 |
| (STZ Ex-9) - CTR SAL | -0.0545 | 0.0331 | 15 | -0.1251 | 0.0162 | -1.6434 | 0.1211 |

| | | | | | | | |
|---|---|---|---|---|---|---|---|
| (STZ Ex-9) - STZ SAL | -0.0014 | 0.0331 | 15 | -0.0721 | 0.0692 | -0.0425 | 0.9667 |
| CTR SAL - STZ SAL | 0.0531 | 0.0313 | 15 | -0.0135 | 0.1197 | 1.6980 | 0.1101 |

Table 20. Ileum TBARS – main effects model

| contrast | estimate | SE | df | lower.CL | upper.CL | t.ratio | p.value |
|---|---|---|---|---|---|---|---|
| CTR - STZ | 8.1165 | 9.1806 | 20 | -11.0339 | 27.2670 | 0.8841 | 0.3872 |
| (Ex-9) - SAL | 20.1114 | 9.1806 | 20 | 0.9610 | 39.2619 | 2.1906 | 0.0405 |

Table 21. Ileum TBARS – interaction model

| contrast | estimate | SE | df | lower.CL | upper.CL | t.ratio | p.value |
|---|---|---|---|---|---|---|---|
| (CTR Ex-9) - (STZ Ex-9) | 26.5062 | 11.4890 | 19 | 2.4595 | 50.5529 | 2.3071 | 0.0325 |
| (CTR Ex-9) - CTR SAL | 40.3400 | 12.0497 | 19 | 15.1197 | 65.5604 | 3.3478 | 0.0034 |
| (CTR Ex-9) - STZ SAL | 28.2279 | 11.4890 | 19 | 4.1812 | 52.2746 | 2.4570 | 0.0238 |
| (STZ Ex-9) - CTR SAL | 13.8339 | 12.0497 | 19 | -11.3865 | 39.0543 | 1.1481 | 0.2652 |

| | | | | | | | |
|---|---|---|---|---|---|---|---|
| (STZ Ex-9) - STZ SAL | 1.7217 | 11.4890 | 19 | -22.3250 | 25.7685 | 0.1499 | 0.8825 |
| CTR SAL - STZ SAL | -12.1121 | 12.0497 | 19 | -37.3325 | 13.1083 | -1.0052 | 0.3274 |

Table 22. Ileum CAT – main effects model

| contrast | ratio | SE | df | lower.CL | upper.CL | t.ratio | p.value |
|---|---|---|---|---|---|---|---|
| CTR / STZ | 1.3681 | 0.3664 | 20 | 0.7826 | 2.3918 | 1.1704 | 0.2556 |
| (Ex-9) / SAL | 1.8305 | 0.4902 | 20 | 1.0470 | 3.2001 | 2.2575 | 0.0353 |

Table 23. Ileum CAT – interaction model

| contrast | ratio | SE | df | lower.CL | upper.CL | t.ratio | p.value |
|---|---|---|---|---|---|---|---|
| (CTR Ex-9) / (STZ Ex-9) | 1.2282 | 0.4642 | 19 | 0.5568 | 2.7090 | 0.5439 | 0.5928 |
| (CTR Ex-9) / CTR SAL | 1.6256 | 0.6444 | 19 | 0.7091 | 3.7267 | 1.2258 | 0.2352 |
| (CTR Ex-9) / STZ SAL | 2.5043 | 0.9465 | 19 | 1.1354 | 5.5236 | 2.4290 | 0.0252 |
| (STZ Ex-9) / CTR SAL | 1.3236 | 0.5246 | 19 | 0.5774 | 3.0342 | 0.7072 | 0.4880 |
| (STZ Ex-9) / STZ SAL | 2.0390 | 0.7706 | 19 | 0.9244 | 4.4973 | 1.8852 | 0.0748 |
| CTR SAL / STZ SAL | 1.5405 | 0.6106 | 19 | 0.6720 | 3.5316 | 1.0902 | 0.2893 |
