## Supplement 2 for "Failure of the brain glucagon-like peptide-1-mediated control of intestinal redox homeostasis in a rat model of sporadic Alzheimer’s disease"

We have observed intracerebroventricular administration in animals anesthetized with ketamine (70mg/kg) and xylazine (7mg/kg) induces significant perturbations in plasma and cerebrospinal fluid glucose concentrations (**Fig S1**). This is rarely discussed in the literature, but such pathophysiological fluctuations that arise as a consequence of inherent methodological limitations might affect experimental results, especially in animal experiments focused on the processes associated with tight metabolic control in the brain or the periphery. No differences between different groups were observed in our experiment, however, the fact that intracerebroventricular administration protocol-induced hyperglycemia acutely generates a pathophysiological rather than physiological cellular milieu should still be emphasized and acknowledged.

Glucose was measured with the standard Trinder method utilizing glucose oxidase and 4-amino-antipyrine provided in a commercial kit (Greiner Diagnostic, Germany). Absorbance was read with Infinite F200 PRO multimodal microplate reader (Tecan, Switzerland).

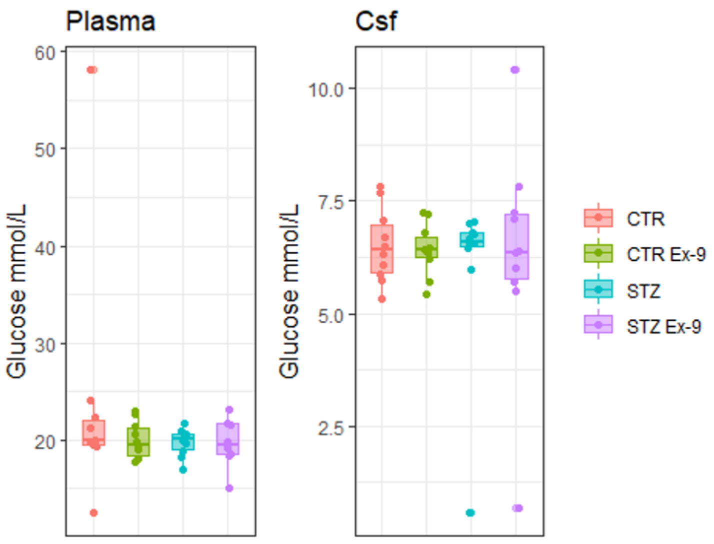

Homolak et al. (2021) Failure of the brain GLP-1-mediated control of intestinal redox homeostasis in a rat model of sporadic Alzheimer’s disease

**Fig S1. Plasma and cerebrospinal fluid (Csf) glucose concentration in the control rats and rats treated with intracerebroventricular streptozotocin (STZ-icv) 30 minutes after intracerebroventricular saline (CTR/STZ) or Exendin3(9-39)amide (CTR Ex-9/STZ Ex-9) administration.**
